# Correlative cryogenic montage electron tomography for comprehensive in-situ whole-cell structural studies

**DOI:** 10.1101/2021.12.31.474669

**Authors:** Jie E. Yang, Matthew R. Larson, Bryan S. Sibert, Joseph Y. Kim, Daniel Parrell, Juan C. Sanchez, Victoria Pappas, Anil Kumar, Kai Cai, Keith Thompson, Elizabeth R. Wright

## Abstract

Imaging large fields of view while preserving high-resolution structural information remains a challenge in low-dose cryo-electron tomography. Here, we present robust tools for montage electron tomography tailored for vitrified specimens. The integration of correlative cryo-fluorescence microscopy, focused-ion beam milling, and micropatterning produces contextual three-dimensional architecture of cells. Montage tilt series may be processed in their entirety or as individual tiles suitable for sub-tomogram averaging, enabling efficient data processing and analysis.

## Main

There is an increasing interest and need for a comprehensive understanding of the structure and function of macromolecules, both isolated and within the context of a larger biological system. While cryo-electron microscopy (cryo-EM) of purified proteins (e.g., single-particle cryo-EM) has propelled the cryo-EM ‘resolution resolution’^1^, *ex-situ* conditions may not capture cellular interactions taking place between biological molecules in a native context. Cryo-electron tomography (cryo-ET) links three-dimensional (3D) contextual visualization and high-resolution structure determination of cryogenically preserved macromolecular complexes in their native cellular environment^2^, unperturbed by purification or chemical fixation and staining^3,4^. Computationally extracted sub-tomograms can be analyzed and classified to reveal sub-nanometer (sub-5 Å) to nanometer (1∼3 nm) resolution structures of *in-situ* complexes^5^. Cryo-ET is generally restricted to investigations of small specimen volumes and the thin peripheral areas of cells (< 500 nm) that are penetrable by the electron beam. To explore thicker regions of cells, sample thinning technologies have evolved and include cryo-electron microscopy of vitreous sections (CEMOVIS)^6^ and cryo-focused ion beam (cryo-FIB) milling^7^. Both methods produce thin sections or lamella from bulk materials, but each may be impacted by preparative artifacts such as sample compression or curtaining, respectively. Cryo-correlative light and electron microscopy (cryo-CLEM)^8^ correlates the temporal and spatial information from fluorescence light microscopy (FLM), with ultrastructural data from cryo-ET of the same region of interest (ROI) while retaining context. In combination with cryo-FIB milling, it is now possible to pinpoint an area of interest deep in the interior of a specimen through 3D correlative cryo-FLM-FIB-ET^9^. However, it remains challenging to bridge the disparity of the spatial scales in multi-modal microscopy pipelines where the field of view (FOV) in wide-field FLM can be 10^5^ times (∼0.1 to 5 mm) that of an EM FOV (∼200 nm to 800 µm)^2,10^.

In cryo-ET, tilt series acquisition of an ROI involves tilting the cryo-preserved specimen along one or two axes^11^ while a series of projection images is incrementally captured on a detector. Tilt series of the same region could be collected at both high and low magnifications for subsequent reconstruction into 3D tomograms to obtain high-resolution structural information^12^ and overall landscape visualization, respectively. However, this could be difficult to implement due to the radiation sensitivity of frozen-hydrated biological materials and the need to use low-dose exposure routines (∼1-3 e^-^/Å^2^/tilt) that must maintain sufficient signal over background noise at each tilted image to support individual frame, image, and tilt series alignment. Advances in detector design have supported the use of larger format detectors for certain applications^13^. Of note, detector size scales exponentially with the volume of the data being digitized, thus imposing challenges to hardware and software infrastructure^14^. These technical hurdles and others^5^ have constrained the application of cryo-ET to fractional volumes of cells that result in significant losses to the FOV and contextual information. An approach for obtaining 3D tomograms that encompass a larger FOV is to collect montages of tilt series. The development of montage cryo-ET has been limited. To our knowledge, only one method^15^ has been reported. As noted by the authors, major challenges included seamlessly joining cryo-EM montage tiles with improved automation^15^, processing large volume stitched tomograms^16^, and sub-tomogram averaging (STA).

Here, we demonstrate our adaptation of the principles of montage tomography, which is routinely applied to resin-embedded samples^17,18^, to frozen-hydrated specimens via montage cryo-ET. This montage cryo-ET data collection routine and automated processing schemes (Supplementary Fig. 1) are robust solutions for the generation of molecular-level resolution 3D reconstructions of vitrified specimens.

We employed image-shift montage acquisition^17,19^ to acquire overlapping tiles of defined size at each tilt increment. Rectangular or square montage tile patterns and tile overlaps were investigated while considering: 1) Fresnel fringes formed inside the image detector FOV from the C2 aperture of an electron microscope^20^ introduce non-uniform illumination and disruption in image signals (Supplementary Fig. 2a). Under a magnification (4.6 Å /pixel) and defocus value (−5 *µm*) typical for cryo-ET collection, the beam fringe artifact was determined to affect up to 3 to 4% of the FOV extending from the outer edge of the 3.11 *µm* illuminated area (Supplementary Fig. 2b) when the outer edge of the beam intersected the camera at ∼4% of its long axis. 2) Under parallel illumination, the beam size relative to the camera frame determines the captured FOV and fringe-unimpacted or ‘fringeless’ area (Supplementary Fig. 2c-d). The FOV affected by Fresnel fringes becomes increasingly evident as the beam size decreases under a constant magnification and defocus (Supplementary Fig. 2d-e). Yet, use of an expanded beam and larger illuminated areas to minimizing in-frame fringes also impacts the sample through excess pre-exposure irradiation (Supplementary Fig. 2c). 3) Robust automation of montage stitching at each tilt requires consistent overlap regions between adjacent tiles. Compared to resin-embedded samples where the image contrast is strong^18^, reliable tile overlaps at a majority of the tilts are essential in vitrified unstained samples to achieve gap-free stitching^15^. 4) Sorting of individual tile tilt series from the complete montage collection for efficient data processing and sub-tomogram averaging is applicable only when each tile frame contains enough fringe-unimpacted FOV and retains the ROI throughout the tilt series.

An inherent limitation of montage cryo-ET is the uneven accumulation of dose in overlapping regions between adjacent tiles, which leads to excessive radiation damage. Therefore, we adopted an additional globally applied translation shift that was calculated from the central tile of a montage pattern (Supplementary Figs. 3, 4, 9a). To quantitatively assess cryo-ET data collection and montage tiling strategies, we developed TomoGrapher, a user-friendly simulation tool to visualize tilt series collection routines and determine global and localized electron dose accumulation (Supplementary Fig. 3). TomoGrapher supports simulation of tilt series acquisition with and without a translational shift extending along spiral paths (Supplementary Fig. 4a-f, Table 1). The shape of the spiral trajectory can be adjusted to vary dose accumulation on each voxel over the full tilt range. Circular beam projections and the associated images gradually elongate to ellipses along the X-axis perpendicular to the tilt axis (Y-axis) as the tilt angle increases (Supplementary Figs. 3b, 4c-d), resulting in stretching of the spiral paths. Optional X-axial correction and cosine-weighting^21^ of dose can be implemented in the simulation process (Supplementary Fig. 3). Within the maximum offset distance permitted to retain the FOV in the full tilt range, simulation results suggest larger translations and axial correction introduce more uniformly spread dose (Supplementary Fig. 4g-h).

Overall, both simulation results and experimental data demonstrate the incorporation of a global translational shift allowed for a more even distribution of the total electron dose across each montage tile and tile overlap regions where accumulated dose would otherwise be much higher (Supplementary Fig. 4, Table 1). We determined that a 15 to 20% overlap in X (fringe-affected axis) and 10% in Y (fringe-unaffected axis) of tile frames was sufficient for automated gap-free tile stitching without human intervention at tilt angles up to ±39° and minimal manual fixation at higher tilt angles (± 40° to 60°) using coordinate-based image cross-correlation^22^ (Supplementary Figs. 5a-c). We explored the addition of rotation to translation of the tile montage pattern to further spread the dose. Our data (Supplementary Fig. 5d) support a common observation that rotation of the in-plane image shift deviates significantly from the designated coordinates as the stage tilt increases^23^. This departure in image shift values introduced by global tile rotation was much larger than the applied translation alone at lower tilts and became particularly irregular as the stage was tilted to degrees greater than ±30°. As a result, the inconsistent overlap between tiles caused problematic stitching, consistent with previous reports^15,18^.

We integrated montage cryo-ET with 2D and 3D correlative cryo-CLEM and cryo-FIB routines using CorRelator^24^, 3DCT^9^, and SerialEM^19^ to image large, targeted FOV along the periphery of HeLa cells (Supplementary Fig. 6) and cryo-FIB-milled lamella near the nucleus of adenocarcinomic human alveolar basal epithelial (A549) cells (Fig. 1, Supplementary Fig. 7). Both cell lines are commonly used for studies of both mitochondrial function and respiratory syncytial virus (RSV) infection^25,26^. A pixel size of 4.6 Å was used to support the large FOV and STA sampling requirements^12^. 2D cryo-CLEM was applied to identify fields of long, filamentous RSV particles budding from metabolically active RSV-infected HeLa cells^26^ for montage cryo-ET data collection (Supplementary Fig. 6). Montage tomograms of RSV particles up to 8 *µm* in length revealed the ultrastructure of intact virions and organization of viral compartments (Supplementary Fig. 6g). To explore mitochondrial function and organization closer to the nucleus in naïve and RSV-infected A549 cells, we coupled 3D targeted cryo-FLM-FIB-milling with montage cryo-ET (Fig. 1, Supplementary Fig. 7). Low toxicity fluorescent nanoparticles (40 nm)^27^ were internalized by the cells and used as FIB-milling “fiducials” to position and adjust milling boxes on-the-fly in the Z and XY planes based on the position of nanoparticles relative to the ROI in the 3D cryo-FLM Z-stack (Supplementary Figs. 7, 8b-c). A square of interest was identified (Fig. 1a) and milling boxes were initially positioned via external markers using 3DCT^9^ and CorRelator^24^ (Fig. 1b, Supplementary Fig. 8c), then further adjusted based on the FIB-milling “fiducials” (Supplementary Fig. 8d-f) during the thinning process. Uniformly-sized nanoparticles can be readily differentiated from electron dense lipid droplets by SEM and cryo-ET (Fig. 1c, Supplementary Fig. 8j-m). Post correlation^28^ of cryo-EM images of lamella with the corresponding pre-milling cryo-FLM section confirmed the locations of ROIs and nanoparticles (Fig. 1d-f, Supplementary Fig. 8h-i). We targeted fluorescently-labeled mitochondria-rich areas near the nucleus where multiple 3×3 montage cryo-ET fields were collected; each montage covering an area of ∼7 × 5.5 *µm* (representative 3×3 montage tile in cyan box, Fig. 1e). The 3D correlative targeting in combination with montage cryo-ET supported the precise acquisition of large FOVs of a FIB-thinned cellular lamella. Within the 3D volume, the arrangement of mitochondria, the Golgi, rough ER, and nuclear pore complex along the nuclear envelope was observed with potential to retrieve high-resolution structural information^12^ (Fig. 1f).

**Fig. 1.**
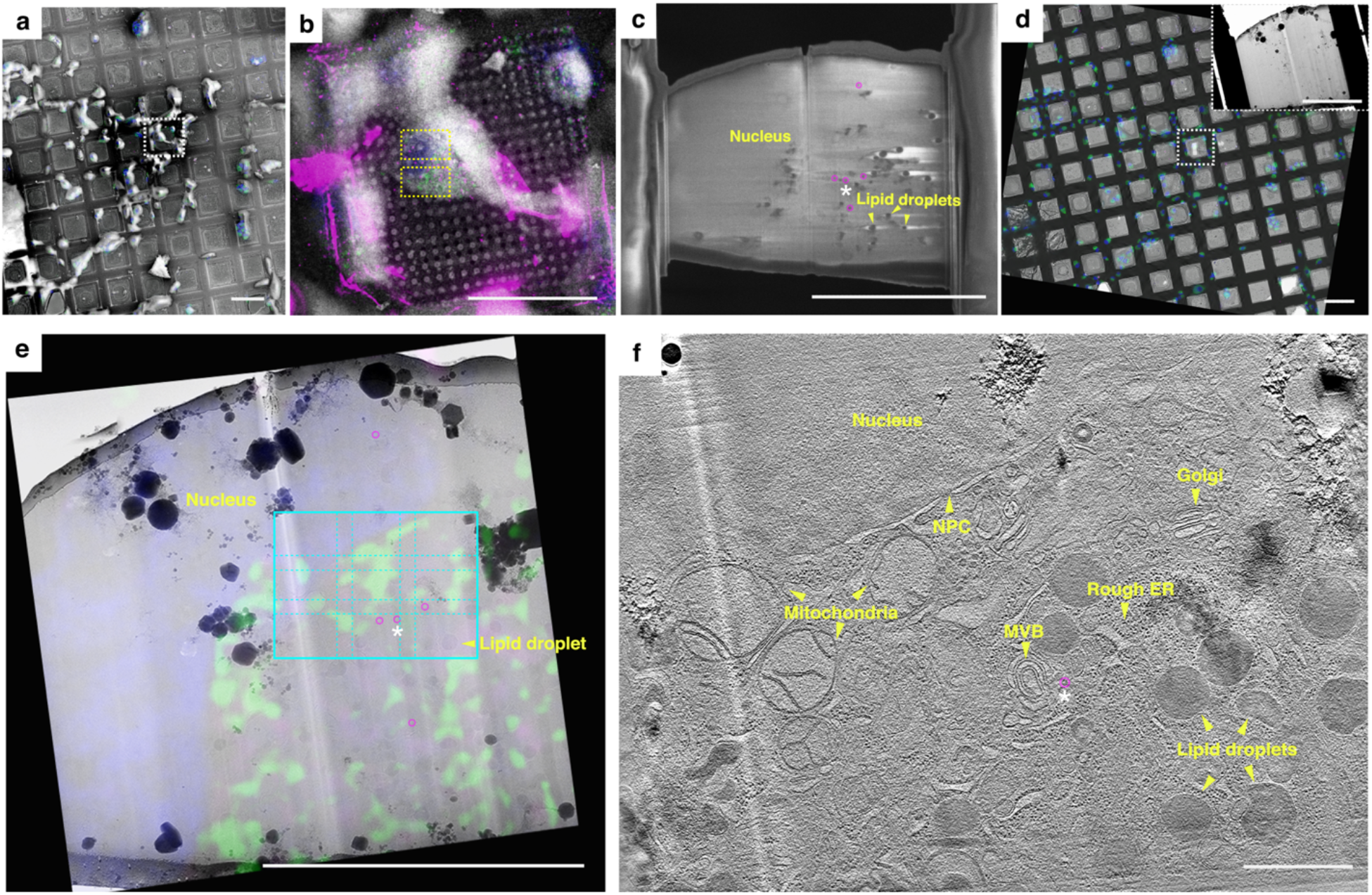
Correlative cryo-FLM, cryo-FIB-SEM, and montage cryo-ET of a cryo-lamella from an A549 cell to visualize the in-situ architecture of the nucleus, mitochondria, and other organelles. **a**. Correlation of cryo-SEM and cryo-FLM grid mosaics to identify an A549 cell in the square of interest (white box). Scale bar, 100 *µm*. **b**. Cryo-FIB-milling boxes (pair of dashed yellow boxes) positioned to target the nucleus (blue) and mitochondria (green) in an A549 cell (white box in **a**), with internalized fluorescent beads (40 nm, pink) as milling “fiducials” for on-the-fly 3D targeted correlation. **c**. Cryo-SEM of the 200 nm thick lamella. Nucleus, lipid droplets, and internalized fluorescent beads (pink circles) are noted. Scale bars, 10 *µm* in **b** and **c. d**. Correlation of pre-FIB-milled cryo-FLM and post-FIB-milled cryo-TEM grid mosaics (Inset, cryo-TEM image of the lamella at higher magnification). Scale bars, 100 *µm* in **d**, 10 *µm* in inset of **d. e**. Post correlation overlay of the 2D cryo-EM image of the lamella with the corresponding Z section from the pre-FIB-milled cryo-FLM stack. A 3×3 tiling for montage cryo-ET (cyan, 6.8 × 5.3 *µm* at a pixel size of 4.603 Å) at the ROI. Scale bar, 10 *µm*. The same internalized fluorescent beads as in **c** (pink circles) are noted. **f**. Tomographic slice, ∼45 nm thick (binned 2x tomogram at a pixel size of 9.206 Å), through the 3×3 montaged cellular tomogram (cyan ROI in **e**). Nucleus, nuclear pore complex (NPC), mitochondria, Golgi apparatus, rough ER, multivesicular bodies (MVB) are noted. Scale bar 1 *µm*. The same internalized fluorescent bead (pink circled) as in **c, e, f** was noted by the asterisk (white).

Next, we applied montage cryo-ET to primary neurons grown on micropatterned cryo-TEM grids^29,30^. Coupling micropatterning with cryo-ET has proven to be valuable for directing cytoskeleton organization and understanding neural outgrowth^30^. Straight-line patterns were used to control neurite growth of primary *Drosophila melanogaster* neurons (Fig. 2a-c). Multiple montage tilt series were collected along neurites protruding from peripheral areas of the cell body (representative 3×4 montage site (∼8 × 7 *µm*) delineated in Fig. 2d). The montage cryo-tomogram revealed the architecture of the cytoskeleton, including continuous microtubules (∼9 *µm*) stretching along the pattern and the presence of actin filaments extending from the neurite. The localization and 3D organization of mitochondria that exhibited matrix regions densely packed with calcium granules was indicative of possible metabolic activities^31^ (Fig. 2e-f). The application of cryo-montage tomography is important for generating large-scale 3D molecular vistas of neurons and other cells that are responding to external physical cues such as those imposed by micropatterning.

**Fig. 2.**
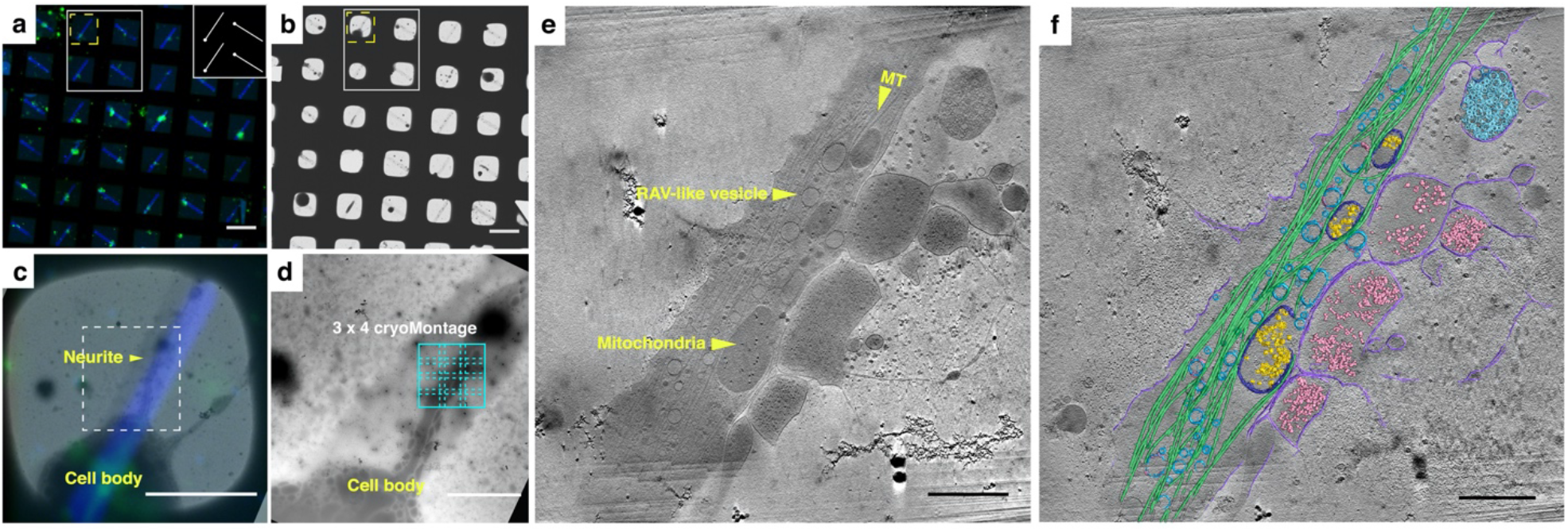
Correlative live-cell-FLM and montage cryo-ET of primary *Drosophila melanogaster* neurons on a mask-free micropatterned cryo-TEM grid. **a-b**. Live-cell FLM (**a**) and cryo-EM (**b**) grid maps of membrane GFP-labeled primary neurons (green) cultured on a gold-mesh, holey (R1.2/20) carbon film micropatterned with a straight-line pattern (inset in **a**) across the grid and coated with fluorescent concanavalin A (blue) to control the growth of neurites. Scale bars, 100 *µm*. **c**. Overlay of correlated FLM-cryo-EM image of the square highlighted in yellow in **a** and **b. d**. Enlarged cryo-EM image of the dashed white boxed region in **c**. A 3×3 tiling for montage cryo-ET (cyan, 6.8 × 5.3 *µm* at a pixel size of 4.603 Å) on the region of interest (ROI) extending from the cell body. Scale bars, 50 *µm* in **c**, 10 *µm* in **d. e-f**. Tomographic slice of ∼45 nm thick (binned 2x tomogram at pixel size of 9.206 Å), reconstructed from the 3×3 montage tilt series at the ROI (**e**) and the corresponding segmentation (**f**). The structured organization of microtubules (MT, green) and the arrangement of surrounding cellular organelles, including mitochondria (dark blue with calcium granules (yellow)), ribosomes (light pink) and ribosome-associated-like vesicles (darker cyan) are noted. Scale bars, 1 *µm*.

A completely reconstructed unbinned tomogram of a fully stitched 3×4 montage tilt series could be ∼700 GB or more, particularly as the sampling pixel size decreases and tile pattern size expands. To maximize output and develop efficient processing schemes, we explored sorting individual tile tilt series and independently reconstructing them into tile tomograms that have the same volume and pixel size as a regular single cryo-tomogram. Consistent with simulation results (Supplementary Figs. 4, 5), each tile tilt series exhibited the same spiral image shift trajectory as the fully-stitched montage tilt series (Supplementary Fig. 9a-b). When the largest image translation offset was restricted to 30% or less of the FOV per image, each tile tilt series maintained the specified ROI within the majority of the tilt series images. After determining the eucentric plane of the montage field, per tilt focus and per group tracking were adjusted using the nearby focus area during the dose-symmetric acquisition. CTF determination using CTFPLOTTER^32^ indicated that the data acquisition scheme provided a relatively stable defocus (± 1 *µm*) over tilt series acquired from a wide range of samples (Supplementary Fig. 9c). We used CTFFIND4 for tilted defocus determination^23^. CTFFIND4 reported the high-resolution limit of detected Thon rings that ranged from 8 to 20 Å for individual tiles and stitched montages at 0° tilt and high tilts (Supplementary Fig. 9d). Per tile defocus adjustment could be implemented to decrease defocus variation. We performed sub-tomogram averaging on the RSV F fusion (F) glycoprotein to compare with our previous results^33^. Viral F glycoproteins are arrayed on the surface of filamentous RSV particles either budding or released from RSV-infected BEAS-2B cells grown on EM grids (Supplementary Fig. 10a-e). Volumes containing F were extracted from individually-reconstructed tile tomograms for STA (n = 23250, ∼29 Å, C3 symmetry imposed). In an effort to determine whether an improved sub-tomogram average would result from the removal of particles located along stitched overlap zones (15% of X and 10% of Y edges), 971 particles from those regions out of the 7947 total unique particles were removed to yield a refined average of F at ∼26 Å resolution (n = 20707, C3 symmetry imposed). These averaged F structures were on par with what has been reported^33^. We anticipate further improvements to the average could be gained, in part, by using a lower defocus range, a larger number of particles, and further refinement of particles included in the average based upon position in individual tile tomograms.

In conclusion, correlative cryo-montage tomography is a workflow suited for capturing both comprehensive fields of view and targeted regions of interest in complex biological environments at molecular-level resolution. The montage cryo-ET tools presented here are applicable for modern TEMs with stable lens systems. Montage cryo-ET is adaptable to existing imaging routines and supports flexible user-defined tile patterns, streamlined data acquisition, pre-processing automation, and maximization of cryo-ET output with both individual tile and montage tilt series. Rectangular array tile montage for cryo-ET has laid the foundation for future developments with super-montage tomography that incorporate both image-shift and stage-shift collection^17^. Cryo-ET tilt series are commonly acquired at the highest tolerable dose for the biological target; future work will assess the impact of total dose, dose distribution, and montage tile stitching on the preservation of high-resolution structural information for both montage and individual tile tomograms. Ultimately, montage tomography solutions will help bridge the resolution gap and field of view losses present between multi-modal microscopy imaging pipelines.

## Acknowledgements

We thank Dr. Jill Wildonger, Dr. Sihui Z. Yang, and Mrs. Josephine W. Mitchell in the Department of Biochemistry, University of Wisconsin, Madison for kindly sharing the elav-Gal4, UAS-CD8 : GFP fly strain (Bloomington stock center, #5146). This work was supported in part by the University of Wisconsin, Madison, the Department of Biochemistry at the University of Wisconsin, Madison, and public health service grants R01 GM114561 and U24 GM139168 to E.R.W. from the NIH. We are grateful for the use of facilities and instrumentation at the Cryo-EM Research Center in the Department of Biochemistry at the University of Wisconsin, Madison.

## Author contributions

J.E.Y. and E.R.W. conceived and designed the study. M.R.L, J.E.Y, E.R.W developed TomoGrapher and pre-processing workflow. J.E.Y., B.S.S, J.Y.K., D.P., J.M.S., V.P., A.K., K.C., A.K., K.T. prepared the samples, performed the experiments, and data processing. J.E.Y. and E.R.W. wrote the manuscript, with contributions from all authors. All authors read and approved the manuscript.

## Methods

### Cell lines and cell culture

HeLa cells (ATCC CCL-2, ATCC, Manassas, VA, USA) and A549 cells (ATCC, CCL-185) were cultured and maintained in supplemented DMEM complete medium and BEAS-2B cells (ATCC, CRL-9609) cultured in supplemented RPMI-1640 complete medium as reported previously^26^. Primary *Drosophila melanogaster* third-instar larval neurons (the strain elaV-Gal4, UAS-CD8::GFP maintained and kindly provided by the Wildonger lab, UCSD) were extracted, cultured in supplemented Schneider’s media, and maintained on micropatterned grids as previously described^34^.

### Cell seeding, infection and *in-situ* labeling on TEM grids

Cell seeding on the TEM grids was performed following previous reports^26^. Briefly, Quantifoil grids (200 mesh Au R2/2 carbon or SiO_2_ film; Quantifoil Micro Tools GmbH, Großlöbichau, Germany) were coated with extra carbon (5 to 8 nm) and glow discharged (10 mA, 60 sec). HeLa and BEAS-2B cells were trypsinized and seeded at a density of 0.7 × 10^5^ cells/mL, followed by an overnight incubation prior to subsequent applications. For montage cryo-ET and correlative cryo-FLM-montage-ET, HeLa and BEAS-2B cells on the grids were infected with the recombinant virus strain RSV rA2-mK^+^ at an optimized multiplicity of infection (M.O.I.) of 10 for 24 hours at 37 °C and 5 % CO_2_, as determined previously^26^. For cryo-focused ion beam milling (FIB-milling), A549 cells were digested and seeded at a density of 0.3 × 10^5^ cells/mL on SiO_2_ Au Quantifoil grids for 16 to 24 hours.

### Micropatterning and neuron cell culture on TEM grids

Micropatterning and culturing of primary *Drosophila* larval neurons was performed as described^30^. Briefly, the extra carbon coated gold Quantifoil grids (200 mesh, R 1.2/20, holey carbon film; Quantifoil Micro Tools GmbH, Großlöbichau, Germany) were glow-discharged, and coated with 0.05% poly-L-Lysine (PLL). The grids were then functionalized by applying a layer of anti-fouling polyethylene glycol-succinimidyl valerate (PEG-SVA), followed by application of a photocatalyst reagent, 4-benzoylbenzyl-trimethylammonium chloride (PLPP) gel. Maskless photopatterning was performed to ablate the anti-fouling layer in defined patterns with a UV laser (λ = 375 nm, at a dose of 30 mJ/mm^2^) using an Alvéole PRIMO micropatterning system. Adherent extracellular matrix (ECM) protein, fluorescently-labeled concanavalin A, Alexa Fluor™ 350 conjugate (emission, λ = 457 nm, 0.5 mg/mL in water or PBS, ThermoFisher Scientific), was then added to promote the cellular adhesion and growth of primary *Drosophila* larval neurons isolated and cultured according to established protocols^34,35^. These neurons had pan-neuronal GFP expression (emission, λ = 525 nm) on the membrane (CD8-GFP) to allow for tracking using live-cell wide-field fluorescent microscopy imaging. Neurons on patterned grids grew for a minimum of 48-72 hours for neurite growth prior to plunge freezing.

### Vitrification

For EM grids prepared for non-FIB cryo-ET applications, 4 µl of 10 nm BSA-treated gold fiducial beads (Aurion Gold Nanoparticles, Electron Microscopy Sciences, PA, USA) were applied before vitrification. The grids were plunge-frozen using either a Gatan CryoPlunge3 system (CP3) with GentleBlot blotters (Gatan, Inc., Pleasanton, CA, USA) or a Leica EM GP (Leica Microsystems). The Gatan CP3 system was operated at 75 ∼ 80 % humidity and a blot time of 4.5 to 5.5 s for double-sided blotting and plunge freezing. The Leica EM GP plunger was set to 25 °C to 37 °C, 99% humidity and blot times of 6 s for R 1.2/20 micropatterned carbon-foil grids, and 15 s for R2/2 SiO_2_ foil grids for single-sided blotting and plunge freezing. Plunge-frozen grids were then clipped and stored in cryo-grid boxes under liquid nitrogen.

### Correlative live-cell and cryogenic fluorescent microscopy

Healthy A549 or RSV-infected HeLa cells were stained with MitoTrackerGreen FM (M7514, ThermoFisher Scientific, 100 nM, 30 min at 37 °C and 5 % CO_2_), washed, and stained with Hoechst-33342 (H3570, ThermoFisher Scientific, 1 to 1000 dilution, 20 min at 37 °C and 5 % CO_2_) to visualize mitochondria and the nucleus. As reported previously^8^, live-cell wide-field imaging (20 X, 0.4NA lens, dry) and cryo-FLM (50 X, ceramic-tipped, 0.9NA lens) on vitrified samples were performed on a Leica DMi8 and Leica EM Cryo-CLEM THUNDER system, respectively. The brightfield and band pass filter cubes of GFP (emission, λ = 525 nm), DAPI (emission, λ = 477 nm), Texas Red (emission, λ = 619 nm), and Y5 (emission, λ = 660 nm) were used. Live-cell wide-field images were collected as a grid montage at 20X. For cryo-FLM, Z-stack projections of 12 to 15 *µm* for each channel were collected on the vitrified sample at a Nyquist sampling step of 350 nm using the Leica LAS X software. Small Volume Computational Clearance (SVCC) from the Leica LAS X THUNDER package was applied for fluorescent image deconvolution and blurring reduction on the cryo-FLM image stacks. All images and mosaics were exported and used as LZW compressed lossless 16-bit TIFF format. The on-the-fly cryo-FLM to cryo-ET correlation and data collection was performed using CorRelator^24^. Grids that were imaged under cryogenic conditions were saved and stored in in cryo-grid boxes under liquid nitrogen.

### 3D targeted Cryo-FIB-SEM

Low toxic nanoparticles of 40 nm (FluoSpheres, carboxylate modified microsphere, dark red fluorescent (660/680 nm), ThermoFisher Scientific, F8789) were introduced to cells seeded on the grid for an incubation of 2 hours at 2 mg/mL, followed by washing with 1X PBS and 5 min incubation of 5 ∼10 % glycerol as a cryoprotectant to properly vitrify cells. Afterwards, 4 µl of diluted 200 nm FluoSpheres (1 to 200 dilution, dark red fluorescent (660/670 nm), ThermoFisher Scientific, F8807) were applied to the grid in the humidity chamber of the Leica EM GP plunger prior to the vitrification step. Following Leica EM Cryo CLEM THUNDER imaging, clipped grids were transferred into a dual-beam (SEM/FIB) Aquilos2 cryo-FIB microscope (ThermoFisher Scientific) operating under cryogenic conditions. To improve the sample conductivity and reduce the curtaining artifacts during FIB milling, the grid was first sputter-coated with platinum (10 mA, 15 to 30 sec), and then coated with organometallic platinum using the in-chamber gas injection system (GIS, 6 sec with a measured deposition rate of 600 nm/sec). A 2D affine transformation on the XY plane was performed to align cryo-FLM and cryo-scanning electron microscopy (SEM) grid mosaics on a rough micron scale and to further correlate square images from two modalities on a fine nanometer scale precision using hole centroids or 200 nm FluoSpheres in CorRelator. The eucentric height of the region of interest on the cryo-shuttle inside the dual-beam microscope and a shallow FIB-milling angle of 8 to 12° were determined. 2D SEM and FIB views of the squares (with the field of view large enough to include sufficient external microspheres and features as registration points) that contained the region of interest (ROI) were collected at the eucentric height and milling angle. 3D coordinate transform between the 3D Z-stack of the Y5 channel (nanoparticles, emission λ = 680 nm) and the 2D FIB view was conducted through the optimized rigid body 3D transformation algorithm available in the 3DCT package using external 200 nm FluoSpheres (n = 4 to 10) as registration points^36^. The transformed coordinates (X, Y, Z) were then imported into CorRelator to fine tune the deviations in X and Y coordinates introduced by the Z transformation in 3DCT, using the closed-form best-fitting least-square solution. The FIB milling boxes were positioned based on the prediction in the 2D FIB view in CorRelator^24^. The ion-beam milling process was performed using 0.3 nA for rough milling and gradually decreased currents of 0.1 nA, 50 pA, 30 pA, and 10 pA, following previously established protocols^37^. Without changing the sample/shuttle position during the milling, a series of cryo-SEM images (electron beam set at 2 kV, 25 pA, dwell time of 1 µs) were collected as the lamella was thinned from an initial thickness of 5 µm, 3 µm, 1 µm, to 800 nm, 500 nm and to the final 200 nm. The SEM images were used to: 1) check the milling process related to stage drift, lamella bending, etc., 2) adjust the milling positions by visualizing the density of internalized 40 nm nanoparticles on the lamella and comparing their positions (X, Y in ∼100 to 200 nm deviation error, and 500 nm in Z relative to the ROI) in the correlated FLM Z-stack, and 3) to confirm the successfully milled isolated region houses the ROI. On-the-fly monitoring of nanoparticle presence provided quick and movement-free feedback on 3D targeted milling when an integrated FLM system is not available. It could also help eliminate excessive alignment steps introduced by shuttle moving in an integrated FLM and FIB-SEM system when the sample is moved back and forth between the FLM imaging and FIB-SEM positions^38^.

To further confirm the preservation of an ROI in the lamella, lamella were transferred to the Leica EM Cryo-CLEM THUNDER system and a Z-stack of the same channels was collected. Post-correlations on lamella of cryo-FLM, cryo-SEM, and cryo-EM were performed using angle-corrected neighboring signals around the lamella to transform cryo-FLM signals to corresponding features on the lamella under cryo-SEM and cryo-EM as described previously^28^.

### Cryo-electron tomography and reconstructions

After live-cell FLM, cryo-FLM imaging, and/or cryo-FIB milling, the same clipped frozen grids were imaged using a Titan Krios (ThermoFisher Scientific, Hillsboro, OR, USA) at 300 kV. Images were acquired on a Gatan Bioquantum GIF-K3 camera in EFTEM mode using a 20 eV slit. Images were captured at various magnifications of 81x (4485 Å/pixel) for whole grid mosaic collection, 470x (399 Å/pixel) and 1950x (177.6 Å/pixel) for square or whole lamella overview, 6500x (27.4 Å/pixel) for intermediate magnification imaging where the field of view is suitable for reliable tracking and 40 nm nanoparticles are visibly distinguished, and 19500x (4.603 Å/pixel) for data acquisition using the SerialEM software package^19^. The full frame size of 5760 × 4092 pixels (counting, CDS mode at 10 eps of dose rate) was collected.

#### Montage cryo-ET setup in SerialEM

Regular image-shift montage acquisition and the multiple record function in SerialEM were adapted for implementation of overlapped beam-image shift tiling. Benchmarks were done at a data acquisition magnification (pixel size of 4.603 Å/pixel) typical for cryo-ET. The illuminated area of 3.1 µm in diameter on the sample was determined by the beam size on the camera and the lens magnification. Fresh gains were collected with this beam size (3.1 µm) and the gain normalized image over vacuum was low pass filtered to 50 Å to enhance the signal of Fresnel fringe peaks. Based on the behaviors of Fresnel fringes^39^ and EM Gaussian signal distribution, the intensity value of the image over pixel was fitted into two distribution curves, a Poisson curve (maximum likelihood estimate/peak *λ*) to fit the edge areas considered as “signal peaks” using 20% of X and Y dimension extending from the edge towards the center, and a Gaussian curve (*µ* ± 2*σ*) to fit the center area considered as noise/background using 90% of whole X and Y dimension from the center towards the edge with a 10% overlap with the “edge” area in MatLab (*poissfit* and *gaussianFit* Curve Fitting Toolbox, MathWorks, Natick, MA, USA). The cut-off from “signal” to “noise” were determined as the possibility of “signal” peaks fading into *µ* ± 2*σ* of “noise” distribution. From multiple measurements (n ≥ 3) along the circular beam edge, the cut-off was 3.5 ∼4 % of X extending from the edge and insignificant in the Y direction. Thus, the rest of the image was considered as a fringeless FOV. Over a wide range of samples, we selected the pixel overlaps of 15% to 20% in the fringe-affected X-direction and 10% in the fringe-free Y-direction as the optimal tiling strategy, such that the usable field of view per camera frame was maximized after correction of Fresnel fringes and optimization of the pre-exposed area outside of camera frame FOV. A regular montage with minimum dimensions of 2×2 was collected with the designated overlaps in X and Y at the data acquisition magnification. A rigorous and reliable image shift calibration in SerialEM at the data acquisition magnification was performed and repeated to ensure a more accurate shift. The beam-image shift tiling information (ImageShift entry under each item section in the image metadata file “.mdoc” file) was obtained on-the-fly. The shift in the X direction to achieve the frame overlap of 15 % in X was retrieved by calculating the difference in image shift between the tile 1 (PieceCoordinates of 0 0 0) and tile 3 (PieceCoordinates of 4608 0 0). The shift in the y direction to achieve the frame overlap of 10 % overlap in Y was retrieved by calculating the difference between tile 1 (PieceCoordinates of 0 0 0) and tile 2 (PieceCoordinates of 0 3516 0). The MultishotParams (X/Y component of image shift vector) was subsequently modified to reflect the tile montage image shift and saved under the SerialEM setting file.

#### Montage cryo-tilt series collection in SerialEM

A SerialEM macro (available at https://github.com/wright-cemrc-projects/cryoet-montage) was used to implement the montage tilt series by acquiring overlapping tiles with designated overlaps to form a montage at each tilt with an additional translational shift and or rotation shift to distribute the dose. Autofocusing was performed at each tilt and shifted along the orientation of the tilt axis 500 nm plus the maximum translational shift of the center of montage tile pattern (e.g. 0.8 µm, Supplementary Figs. 9a) away from the edge of the montage tile pattern. The total dose per tile tilt series was 30-40% below the maximum dose the sample was able to tolerate before evidence of punctate bubbles. At the beginning of each grouped tilt, high magnification/data acquisition tracking with a threshold of 5% of the FOV to acquire a new tracking reference and to iterate the alignment until the threshold was met (usually within one or two iterations), using the nearby autofocusing area, was performed. An additional lower intermediate magnification tracking on a larger FOV of the region of interest was implemented when the tilt angles were above 30°, with a threshold of a difference greater than 10% (usually within one or two iterations). The tilt series collection was paused when the iterations for convergence exceeded 5 times. Benchmarks were done using a 3×3 or 3×4 montage tile pattern and a dose-symmetric scheme running at ± 51° with 3° increments, groups of 3 tilts (original “Hagen scheme” is group of 1 tilt) and a dose of 2 e^-^/Å^2^/tile per tilt for RSV-infected BEAS-2B and HeLa cells, and neurons on micropatterned grids, while a dose of 1.42 e^-^/Å^2^/tile per tilt for A549 cell lamella based on the dose tolerance measurements of each sample. The CDS counting mode and dose rate of 10 eps were used on a K3 camera. The translational shift off-sets followed the global spiral displacement (A_final_ = 1.5, Period = 3, Turns = 50, Revolutions = 15 as input parameters to control the spiral size and resulting displacement offsets). The speed of collection varied with the size of the montage tile, the type of microscope, detector used and sample-dependent total dose. Benchmark collections were 60 min on average and rendered 9 sub-tilt series (3×3 tile pattern) that were stitched seamlessly to form one montaged tilt series. Translational spiral off-sets and collection schemes (bidirectional, dose-symmetric, defocus, etc.) can be adjusted accordingly in the SerialEM macro. The SerialEM macro can be modified to adjust translational spiral off-sets and collection schemes (bidirectional, dose-symmetric, defocus, etc.). The macro can be integrated into existing common automated data collection schemes using the function *Navigator Acquire at points* in SerialEM^40^.

#### Montage tilt series processing

All movie frames per acquisition (0.1∼0.2 e^-^/Å^2^/frame) were aligned and corrected using MotionCor2^41^. For montage tilt series, the total tiles per tilt were registered and stitched together using the designated beam coordinates supplied to the microscope as described above and with linear cross-correlation methods^42^. Despite the fringes on the edge, intrinsic low contrast and low dose received by cryo-tilt series, the regularity of the montage tiling pattern and sufficient overlaps with optimized tiling strategy consistently provided robust and automated seamless stitching without user intervention up to ± 39°. Manual image alignment (MIDAS), implemented in the IMOD package^42^, was used to adjust piece coordinates and iteratively cross-correlate adjacent tiles at higher tilts when necessary. Fully stitched montage tilt series were binned by 2. Tilt series alignment and tomographic reconstructions were performed using the IMOD package with a final pixel size of 9.206 Å. In the absence of gold fiducials in the FIB-milled lamella, alignment of the tilt series was performed using patch tracking or internalized nanoparticles as tracking markers. CTF correction using ctfplotter and ctfphaseflip and dose-weighted filtering^43^ were applied to the aligned tilt series prior to tomogram reconstruction. For segmentation, the aligned tilt series were further binned 3X (final pixel size of 27.6 Å, binned 6X) prior to the tomogram reconstruction. Tomograms were either processed using fast edge-enhancing denoising algorithm based on anisotropic nonlinear diffusion implemented in TomoEED^44^ or Gaussian low-pass filtered to 80 Å for visualization and segmentation. Tomograms of stitched montage tilt series (final pixel size of 27.6 Å) were annotated using convolutional neural networks implemented in the EMAN2 package^45^. For the sub-tilt series, motion-corrected frames were sorted to generate individual tile tilt series that were CTF estimated using CTFFIND4^46^. Tilt series that contained one or more inadequate projections (not properly tracked or failed CTF estimation) were discarded. Qualified sub-tilt series were then aligned, CTF corrected with ctfplotter and ctfphaseflip, dose-weighted filtered, binned to 2X (a final pixel size of 9.206 Å), and reconstructed into tomograms, similar to stitched montage tilt series implemented in the IMOD package. Python or bash scripts (available at https://github.com/wright-cemrc-projects/cryoet-montage) were used to automate the movie frame alignment, montage tile stitching, sub-tilt and montage tilt generation.

### Sub-tomogram averaging

For proof of concept, all averaging was done on unfiltered sub-tilt tomograms using PEET 1.15.0^47^, following the steps reported previously^33^. Particles were manually picked from low pass filtered sub-tilt tomograms binned 2X to a final pixel size of 9.206 Å. Initial particle orientation and rotation axes of particles were generated using SpikeInit (PEET). An initial alignment was done with 565 particles from one tomogram using post-fusion F glycoprotein (EMDB-2393)^48^ low pass filtered to 60 Å as the initial reference. The final average from this first run was low pass filtered to 60 Å and used as the initial reference for a second run on 12,435 particles from a total of 12 sub-tilt tomograms, with a soft-edged cylinder mask applied during the alignment. Duplicate particles were removed during each iteration with a tolerance of 73.6 Å (8 voxels). An additional iteration was done with refined particles using a smaller search range and a larger mask with softer edges. The final average from the second run suggested a three-fold symmetry, consistent with reported crystal structures^49^. C3 symmetry was imposed using *modifyMotiveList* in PEET to generate a three-fold symmetric data set. The C3 symmetric data set was aligned and averaged using the final average from the second run, low-pass filtered to 60 Å as the initial reference for refinement with a smaller search range. The final sub-tomogram average (final pixel size of 9.26 Å, binned 2X) from the C3 symmetry expanded particles was reconstructed from 23,259 particles. The final density map of F was low pass filtered to 29.53 Å based on the FSC cutoff of 0.5 that was calculated in PEET. The picked particles that were in the stitched area (15% in X and 10% in Y) were removed and the rest were reprocessed following the same steps. The final sub-tomogram average (final pixel size of 9.26 Å, binned 2X) from the C3 symmetry expanded particles was reconstructed from 20,707 particles. The final density map of F was low pass filtered to 27 Å based on the FSC cutoff of 0.5 calculated in PEET. The atomic crystal model of pre-fusion F trimer (PDB: 4JHW)^49^ was fitted in the filtered electron density map in Chimera^50^.

### TomoGrapher development

Simulations of the tiled montage imaging and electron beam exposures on a sample were developed as a collection of C# classes built on the Unity 3D engine (version 2020.3.20f1). The simulation represents an array of M x N x O volumes called voxels as a sample interacted by an electron beam. The canvas “stage” comprises of 150 × 150 voxels (X = Y = 150, Z = 1, total extent of 10 × 10 × 0.2 µm) with the center butterfly ROI spanning across 5 × 4 µm on the imaging XY plane. An interactive GUI describes parameters of the imaging including sampling pixel spacing, illuminated area of the beam on the camera, the tiling pattern of the beam, tilt ranges, and translations defined by spiral offsets. A complete run of a simulation iterates through each tilt increment across the range, and ray traces from a sampling position of the illuminated area in the direction of the beam to find intersections on the stage of voxel volumes. Voxels intersected by the rays are aggregated in a set, and each voxel in the set has its total dose incremented once per beam. The viewer provides a real-time animation of the beam shifts, tilting of the stage, and overall exposure at each voxel. TomoGrapher release versions and source code are available at https://github.com/wright-cemrc-projects/cryoet-montage.

### Statistics and Reproducibility

Experiments performed in Figs. 1b-e, 2a-b, Supplementary Figs. 2, 3b-c, 6, 8, 9 were performed independently three times with similar results. Experiments from Figs. 1f, 2c-f, were performed in duplicates, resulting in 7 montaged tilt series on FIB lamellas from two lamellas on two different grids and 13 montaged tilt series on two different micropatterned neuron grids.

## Supplementary Figures and Legends

**Supplementary Figure 1.**
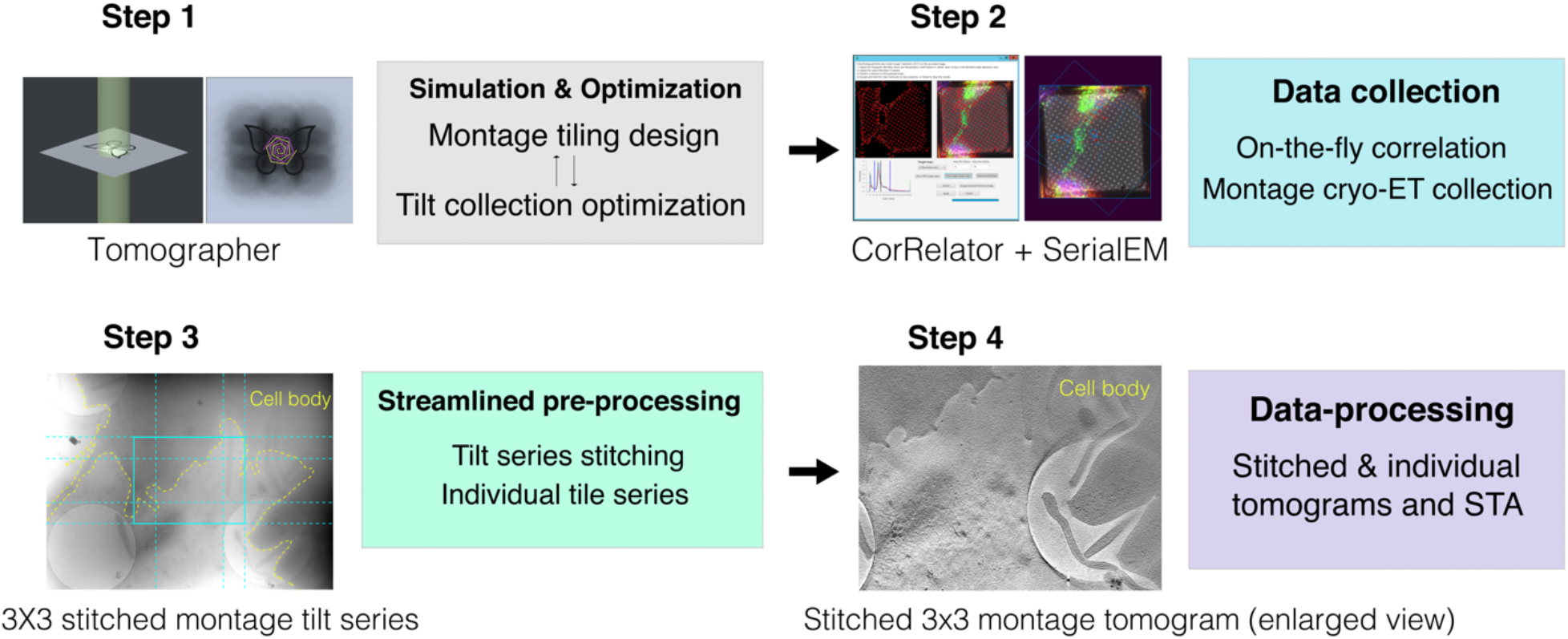
Correlative montage cryo-electron tomography workflow. The correlative montage cryo-electron tomography (cryo-ET) workflow follows four main steps: **Step 1**. *TomoGrapher* simulation-guided tiling design and optimization of tiling patterns for cryo-ET data collection. **Step 2**. CorRelator-driven 2D and 3D cryo-correlation for high-accuracy targeting to support cryogenic tilt series collection with SerialEM. **Step 3**. Automated pre-processing to sort individual tiles and sub-tilt series, and stitch full montages concurrently with data collection. **Step 4**. Analyses including sub-tomogram averaging (STA) and segmentation may be conducted on both individual and stitched montage tomograms.

**Supplementary Figure 2.**
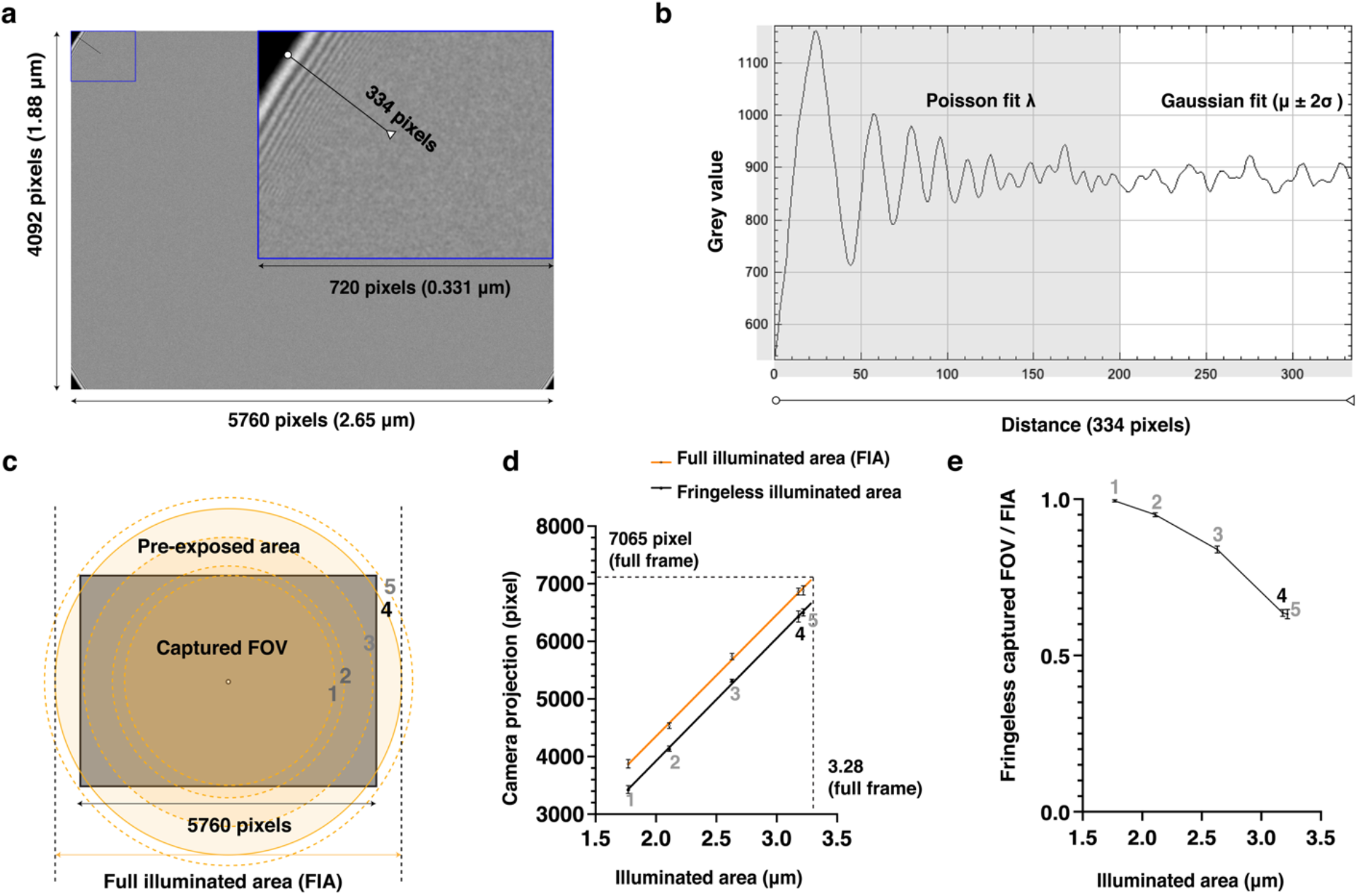
Optimization of beam size and tiling strategy on a triple-condenser lens transmission electron microscope. Benchmarking of the montage cryo-ET workflow was done on a standard Titan Krios 300 kV microscope system with a K3 camera (5760 × 4092 pixels) at a pixel size of 4.603 Å, C2 aperture of 100 mm, and defocus of -5 *µm*. **a**. Fresnel fringes extending over the illuminated area (beam size 3.11 *µm*). ***Inset:*** Enlargement of the boxed fringe-containing area, low pass filtered to 50 Å. **b**. Radial intensity profile along the black line in the ***Inset*** of **(a)**. The region visibly impacted by the Fresnel fringes was determined to extend from the beam edge to ∼200 pixels (grey area) towards the center of the image, affecting between 3.5 to 4 % of the outer edge of a tile. Based on the Fresnel fringes’ Poisson behavior and TEM signal’s gaussian distribution, the fringe peak intensities were fit to a Poisson curve (maximum likelihood estimate/peak *λ*), considered as “signal” and the fringeless illuminated area was fit to a gaussian function (*µ* ± 2*σ*) as “noise” in MatLab (*poissfit* and *gaussianFit* Curve Fitting Toolbox). The cut-off from “signal” to “noise” were determined as the possibility of “signal” peaks fading into *µ* ± 2*σ* of “noise” distribution, from multiple measurements (n ≥ 3) along the circular beam edge. **c-e**. Optimization of beam size (1 to 5, beam sizes increase) in consideration of the full illuminated area (FIA), fringeless illuminated area, pre-exposed area, and captured field of view (FOV) at the constant benchmark magnification and defocus. **c**. The camera FOV (rectangle) and projected beam size (circles) extending outside viewable area. **d**. Linear relationships between illuminated areas on a triple-condenser lens Titan Krios system, defined as the electron beam projected onto the sample, and the beam projected size on the camera plane (orange line, orange circles in **c**, *r2* = 0.9989), and the associated fringeless illuminated area (black line, *r2* = 0.9995). **e**. The ratio of fringeless FOV captured on the camera over the full illuminated area (FIA). Overall, beam size 4 (3.11 *µm*) achieved reasonable fringeless illumination without wasting electron dose on unimaged yet illuminated sample area, and used for benchmark montage cryo-ET in the paper.

**Supplementary Figure 3.**
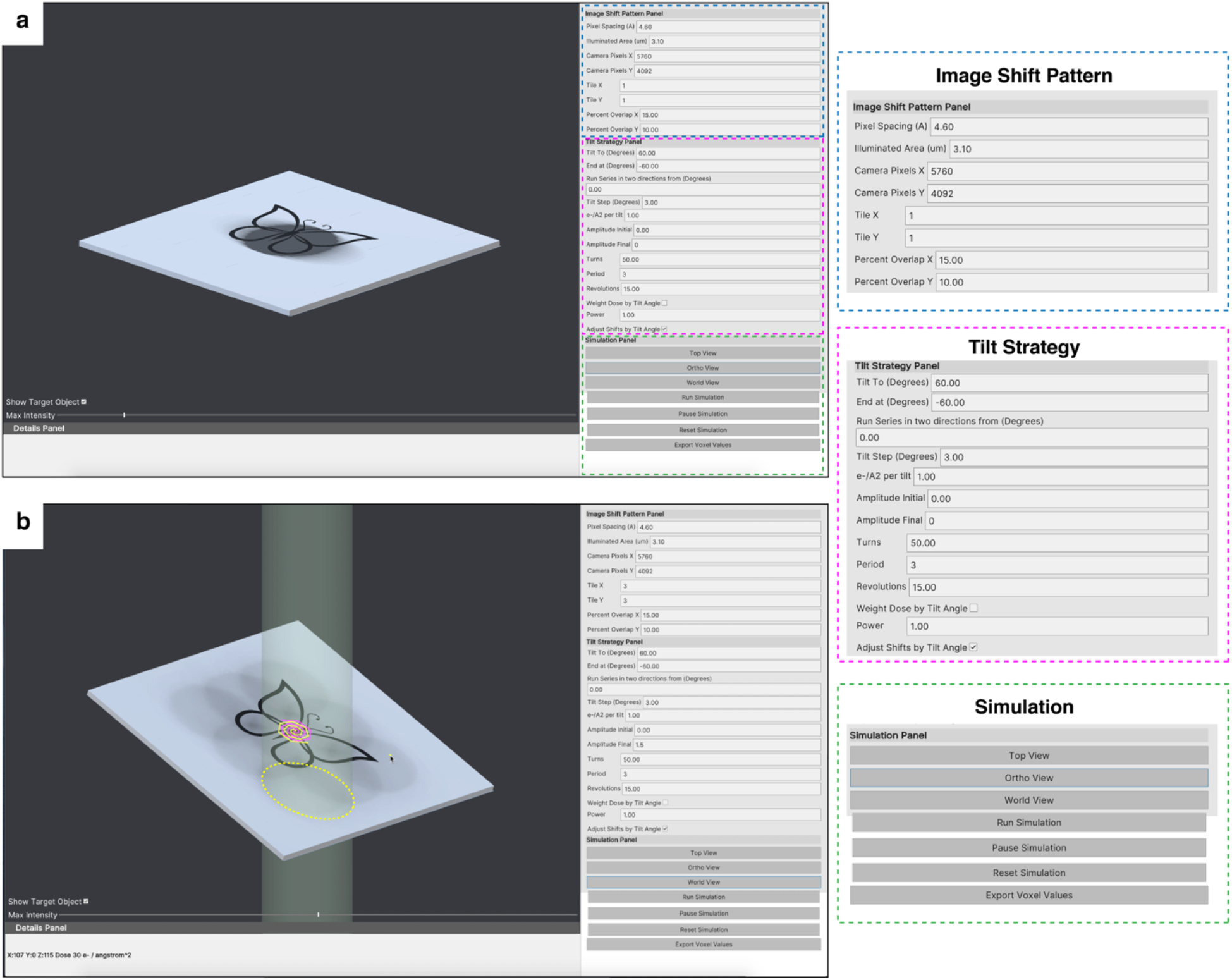
Simulation of tilt series collection in TomoGrapher GUI. *TomoGrapher* is a simulation tool providing a full 3D visualization of **(a)** single-shot and **(b)** montaged (e.g., 3×3 tile) tilt series collection. A stage comprised of 150 × 150 × 1 voxels with a target region of interest (ROI, butterfly, butterfly, 5 *µm* x 4 *µm* x 0.2 *µm*) shows the beam position and accumulated dose values at each position of the total exposed area. The “Image Shift Pattern Panel” (blue boxed) provides parameters to adjust the pixel size, illuminated area (beam size), camera dimensions, and number of and percent overlap for the montage tiles. The “Tilt Strategy Panel” (magenta boxed) controls the tilt range, directionality, tilt step increment, dose per tilt, the amplitudes and geometries of the spiral translational shifts during a tilt series, and provides updated preview images of simulation parameters. Varying intensity as 1/cosine of tilt angle to a certain power is a selectable option to *Weight Dose by Tilt Angle*. Scaling the X-axis shift offset by the cosine of the tilt angle can also be enabled with *Adjust Shifts by Tilt Angle* where the Y-axis is aligned as the tilt axis. The “Simulation Panel” (green boxed) has buttons to change the viewing angle, start a simulation, and export data.

**Supplementary Figure 4.**
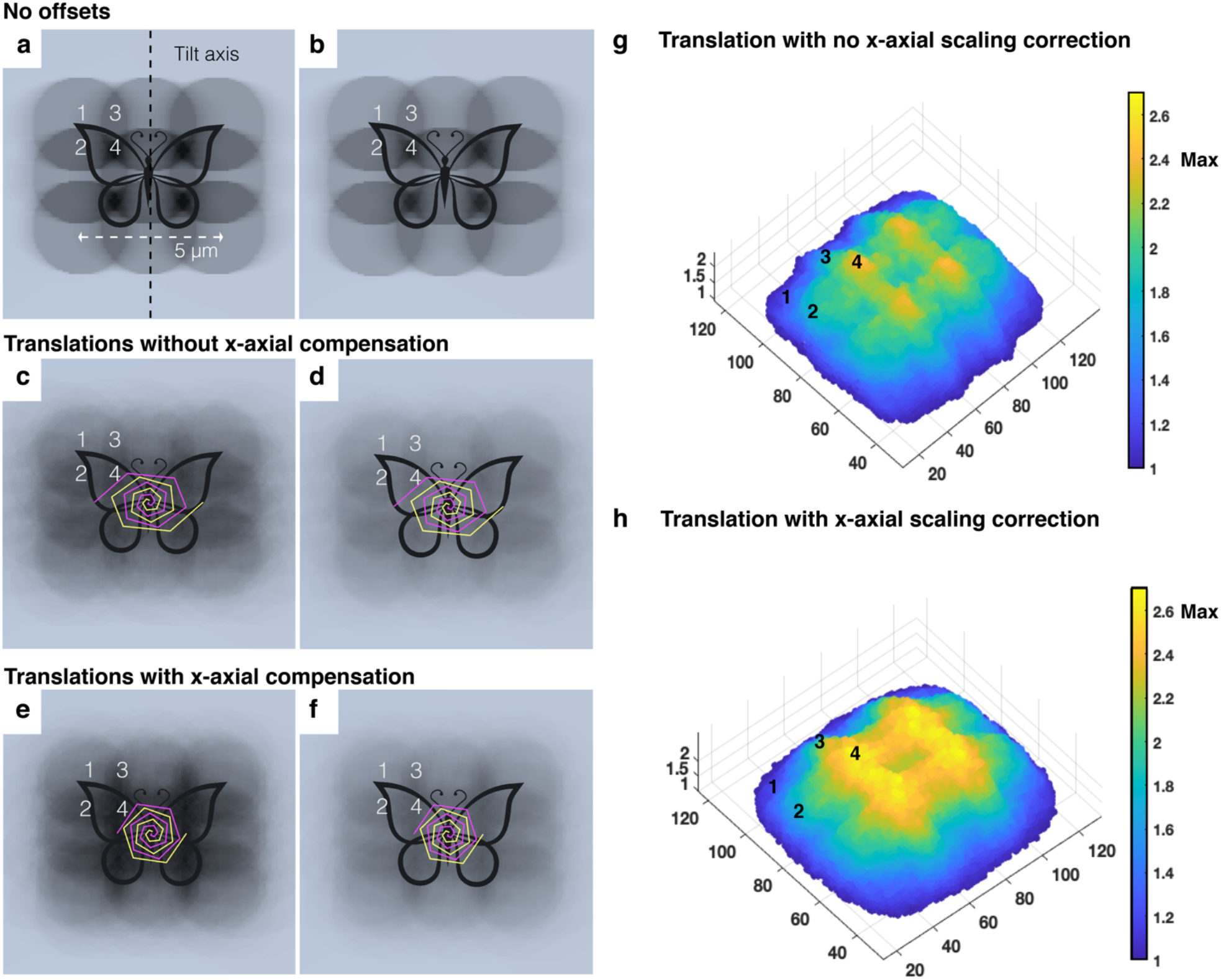
Optimization of montage tiling translational shift for dose distribution in TomoGrapher. Simulated spatial distribution of dose accumulation in six montage tilt series collection schemes (3×3 tile pattern, pixel size of 4.603 Å, beam size of 3.1 *µm*, from - 51° to 51° tilts with a 3° increment, the butterfly as ROI), with no translational offsets **(a, b)**, spiral translational offsets **(c, d)**, and the spiral translational shift with an optional scaling factor (cosine of tilt angle) to reduce *x*-axis shifts at higher tilt angles **(e, f)**. Values provided for dose are 1 e^-^/Å^2^ per tile per tilt in **(a, c, d)** and 0.7 e^-^/Å^2^ per tile per tilt in **(b, d, f)**. The accumulated dose at four areas (10 voxels pivoting around the central 1 to 4 points) were compared between the six collection schemes, detailed in **Supplementary Table 1**. The spiral shift pattern is delineated in pink (0° to 51°) and yellow (−51° to 0°) in **(c-f)** where the same global translational offset is applied (A_final_ = 2.5, Period = 3, Turns = 50, Revolutions = 15). **g-h**. Visualization of the dose distribution received by voxels of the ROI butterfly during the tilt series schemes applied in the benchmarking experiments (A_final_ = 1.5, Period = 3, Turns = 50, Revolutions = 15). The simulated dose values (e/Å^2^/voxel) were exported and plotted in MatLab to show the normalized distribution against the average dose of corresponding tiles from the no translational shift scheme. The same 1-4 center points are marked. The maximum (Max) normalized change is ∼2.4-fold in **(g)** and ∼2.6-fold in **(h)** with the axial scaling compensation. Height and color of each voxel correspond to its accumulated dose. No axial scaling compensation introduces larger shifts and more dose distribution due to the elliptical projection, but the scaling compensation distributes the dose more evenly.

**Supplementary Figure 5.**
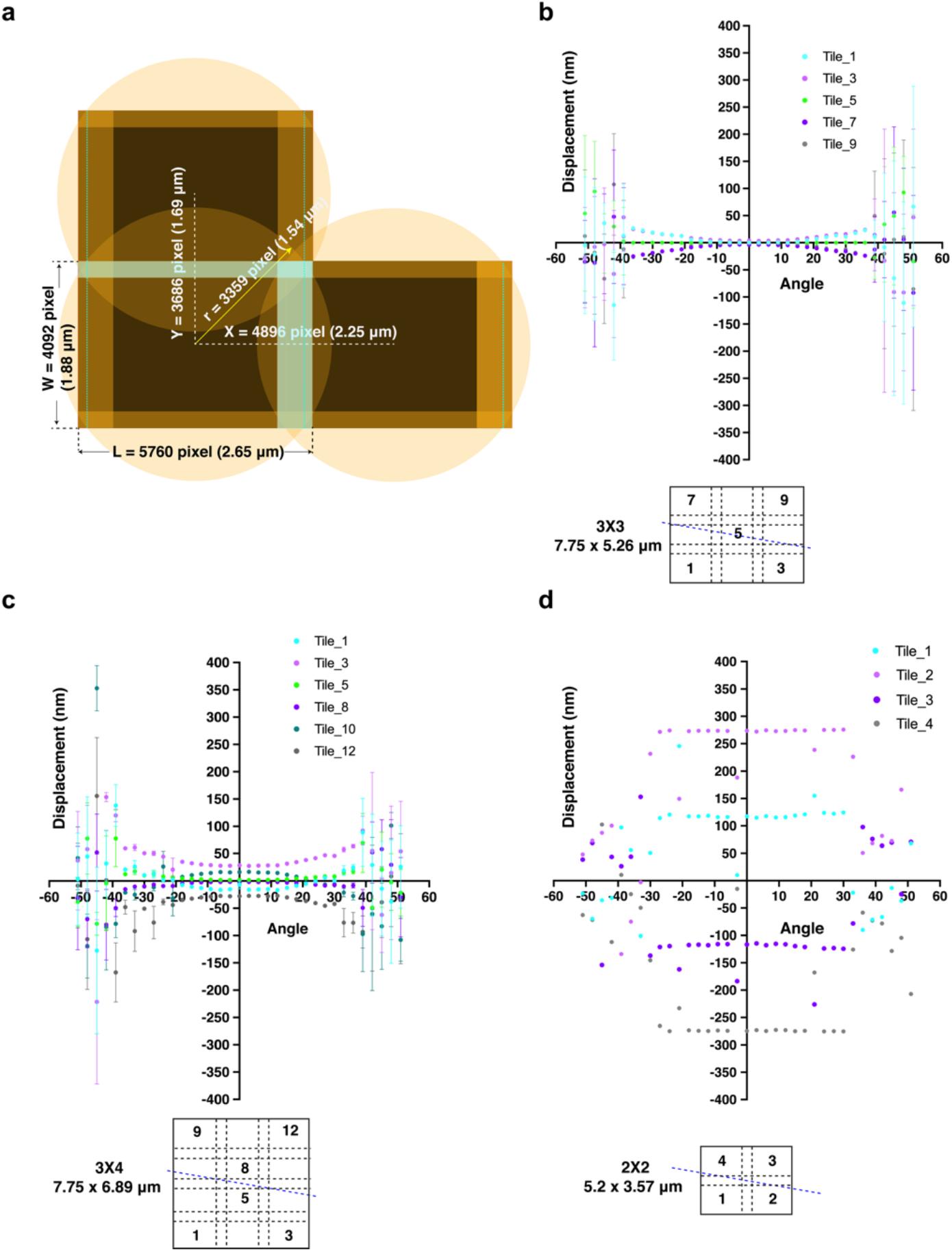
Optimization of the tiling montage cryo-ET tilt series collection with global shift offsets to distribute the electron dose. **a**. Strategy for placement of beam tiles (fringeless usable camera FOV / FIA = 0.7) with 15% to 20% in X (Fresnel fringe-affected) and 10% in Y in a square or rectangular array after balancing fringes, pre-exposed area and the ratio of fringeless captured FOV over FIA. Higher overlap in X supports gap-free stitching under low-dose conditions. **b-d**. Shift displacement of individual tiles (blue dotted line notes the tilt axis) of tilt series collected using the benchmark imaging condition on a Titan Krios 300kV (pixel size of 4.603 Å, C2 aperture of 100 mm, defocus of -5 *µm*, tilt range of -51° to 51°, 3° increment, dose symmetric scheme with group of 3 tilt angles per switch) with application of an additional translation-only (3×3 in **b** and 3×4 in **c**, maximum global translational offsets of 0.8 *µm*, n = 8) or translation plus an additional 20° per tilt rotation offset (total rotation of 35 tilts x 20° = 700°) in (**d**, n = 1) for dose distribution. **d**. Representative plot of tile displacement for a montage tilt series including both translational and rotational shifts which introduce much larger displacements that reduce the effectiveness of automated montage stitching routines. The benchmark spiral parameters (A_final_ = 1.5, Period = 3, Turns = 50, Revolutions = 15) resulted in a ∼0.8 *µm* maximum translational offset.

**Supplementary Figure 6.**
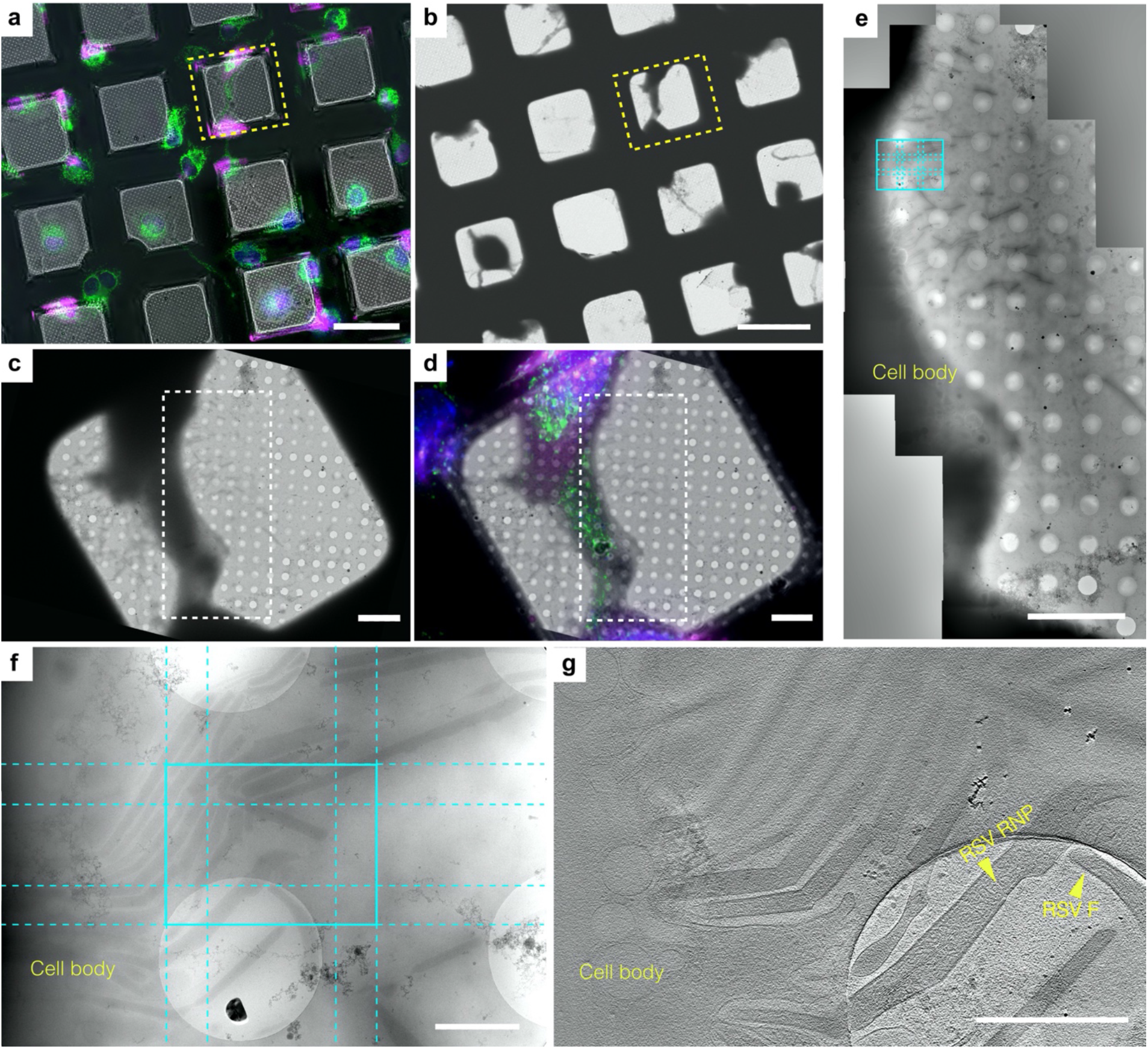
Large field of view of budding RSV virions captured by a correlative cryo-FLM montage cryo-ET workflow. Cryo-FLM **(a)** and cryo-EM **(b)** grid image montages of RSV-infected HeLa cells (magenta) grown on a SiO_2_ filmed R2/2 gold-mesh grid. Mitochondria (green) and nucleus (blue) are labeled. Square of interest highlighted in yellow. Scale bars, 100 *µm*. **c**. Magnified cryo-EM view of the yellow square in **(b). d**. On-the-fly CorRelator-based cryo-FLM and cryo-EM 2D correlation (yellow square in **a, b**) for the ROI identification where montage tilt series were collected in SerialEM. Scale bars, 10 *µm*. **e**. Magnified cryo-EM image montage of the white boxed area in **(d)**. A 3×3 tiling for montage cryo-ET at the ROI (cyan tile pattern) where mitochondrion accumulate and RSV actively bud from the cell plasma membrane. Scale bars, 10 *µm*. **f**. Montage cryo-ET tiles at 0° tilt (cyan, 6.8 × 5.3 *µm* at pixel size of 4.603 Å). Scale bar, 1 *µm*. **g**. Tomographic slice, ∼20 nm thick, through the 3×3 montaged tomogram (cyan ROI in **e, f**). RSV particles with clearly resolved fusion F glycoproteins and ribonucleoprotein (RNP) complexes are present at budding sites along the cell body. Scale bar, 1 *µm*.

**Supplementary Figure 7.**
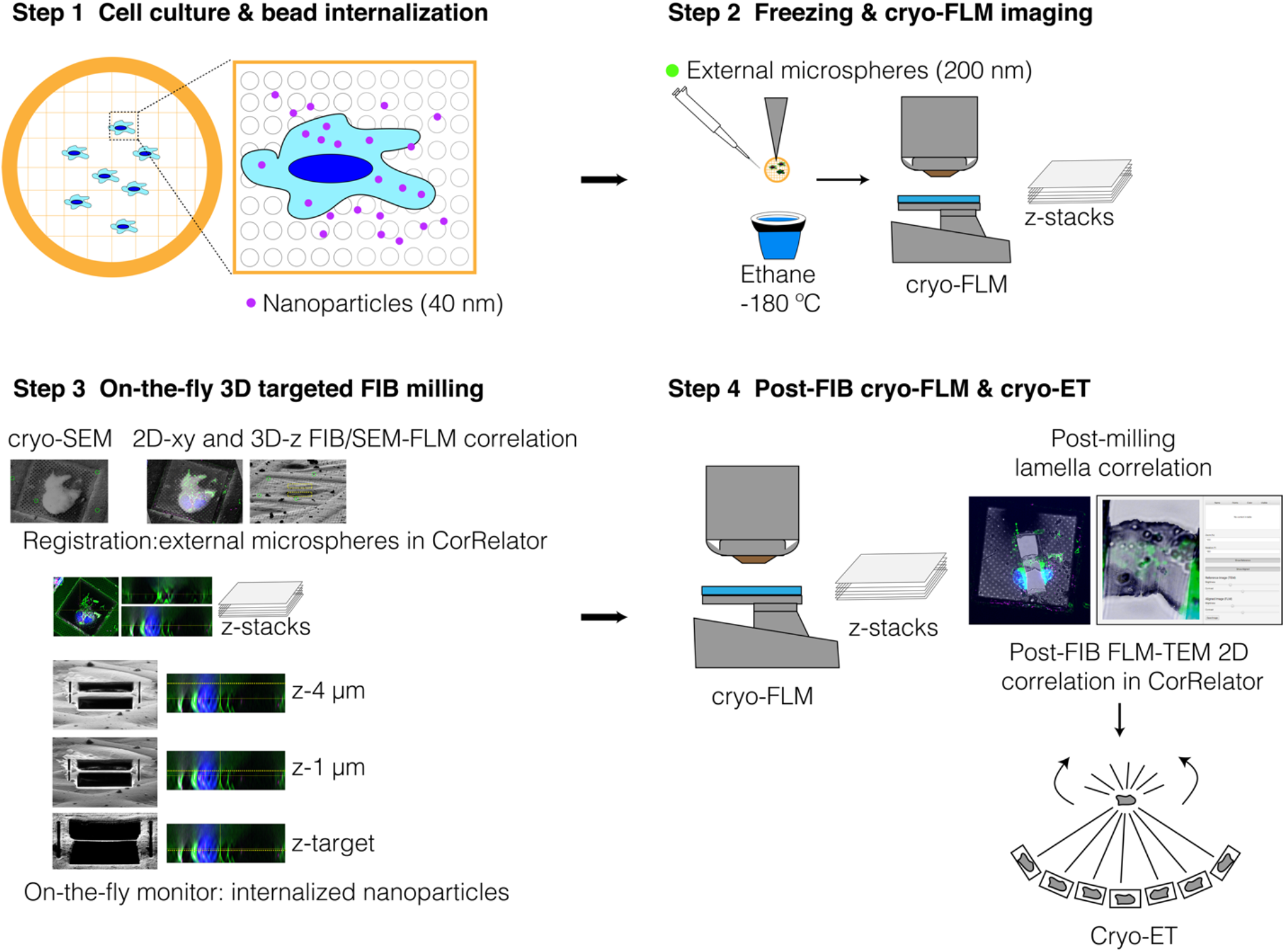
Correlative 3D cryo-FLM-FIB-ET workflow. Overview of the correlative 3D cryo-FLM-FIB-ET workflow using two fiducial markers: 200 nm external fluorescent microspheres for coarse alignment and registration and cell-internalized 40 nm nanoparticles for on-the-fly 3D correlation during cryo-FIB milling. **Step 1**. Fluorescent nanoparticles (40 nm, pink) incubated with cultured cells were internalized. **Step 2**. Fluorescent microspheres (200 nm) were added prior to plunge freezing and grids were then imaged under a cryo-FLM system to acquire Z-stacks of regions of interest (ROIs). **Step 3**. The same grid was loaded onto and imaged with a dual-beam cryo-FIB-SEM system. Using the 3DCT Toolkit and CorRelator, cryo-FLM Z-stacks and 2D cryo-SEM and cryo-FIB images of ROIs were correlated for milling using 200-nm microspheres (FIB view, green circles) in X, Y, and Z. The placement of the milling boxes was refined based on the relative positions of the nanoparticles (pink) and signal of interest (green). **Step 4**. The same FIB-milled lamella was returned to the cryo-FLM system to acquire a new Z-stack cryo-FLM images for post-FIB-milling correlation. The lamella was then loaded into a cryo-TEM system. CorRelator-directed correlation between cryo-FLM and TEM images of the lamella was performed, and montage or regular tilt series were collected with SerialEM.

**Supplementary Figure 8.**
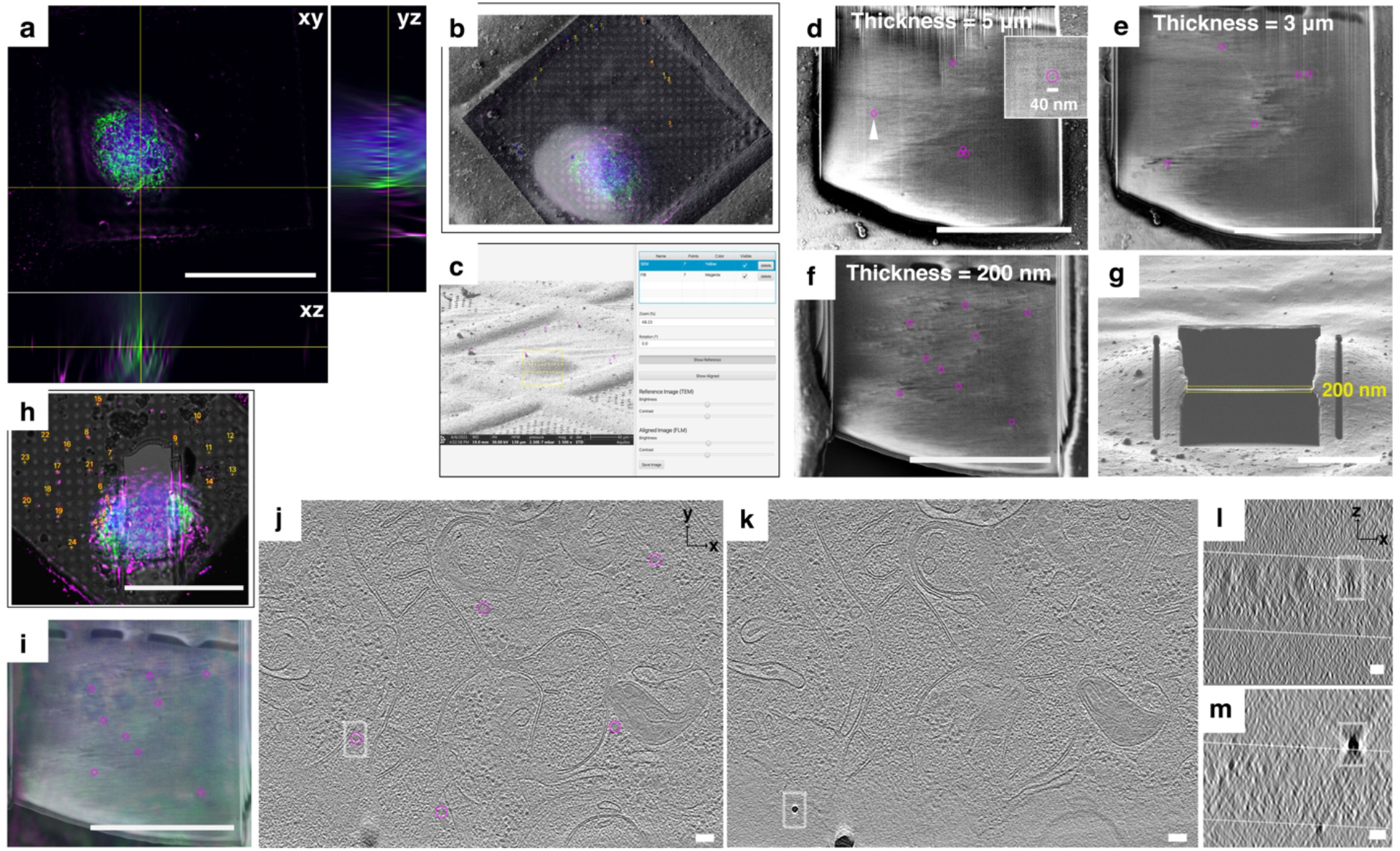
3D correlation using internalized low-toxic fluorescent beads as fiducial markers in cryo-FIB-SEM. a. Orthogonal merged deconvoluted fluorescent cryo-FLM (NA = 0.96) projections of labeled mitochondria (green), nucleus (blue), and internalized nanoparticle (pink) inside the HeLa cell. Scale bar, 100 *µm*. **b-h**. On-the-fly 3D targeted FIB-milling delineated in the Step 3 in Supplementary Figure 6. The X-Y plane correlation is done in CorRelator **(b)** and initial X-Z correlation to position milling boxes in Z (yellow boxes) done in 3DCT and CorRelator **(c)**. Nanoparticle intensity visible under cryo-SEM as the lamella milling progresses from a thickness of 5 *µm* **(d)** to 3 *µm* **(e)**, to the final 200 nm **(f, g)**, highlighted by pink circles (40 nm in diameter). A representative nanoparticle in **(d)** was pointed out by white triangle and enlarged in the inset. **h-i**. 3D cryo-FLM post-correlation of the post-FIB lamella and pre-FIB corresponding fluorescent section in CorRelator. The corresponding nanoparticles in **(f)** are highlighted in pink circles. Scale bars, 10 *µm* in **(d-g)** and **(i)**, 50 *µm* in **(h)**. Red and yellow numbers in the GUI screenshots of **(b, c, h)** are registration points used in CorRelator. **j-m**. Representative X-Y tomographic slices (thickness of 20 nm, pixel size of 4.603 Å) displaying the nanoparticles (pink circle, **j**) and similar sized non-nanoparticle intensity (white boxed, **k**), and their corresponding X-Z projections **(l-m)** of white boxed area in **(j)** and **(k)**. The tomogram Z-volume boundaries are indicated by white dashed lines in **(l-m)**. Scale bars, 50 nm in **(j-m)**. Nanoparticles are less electron dense than ice particles or regular gold fiducials.

**Supplementary Figure 9.**
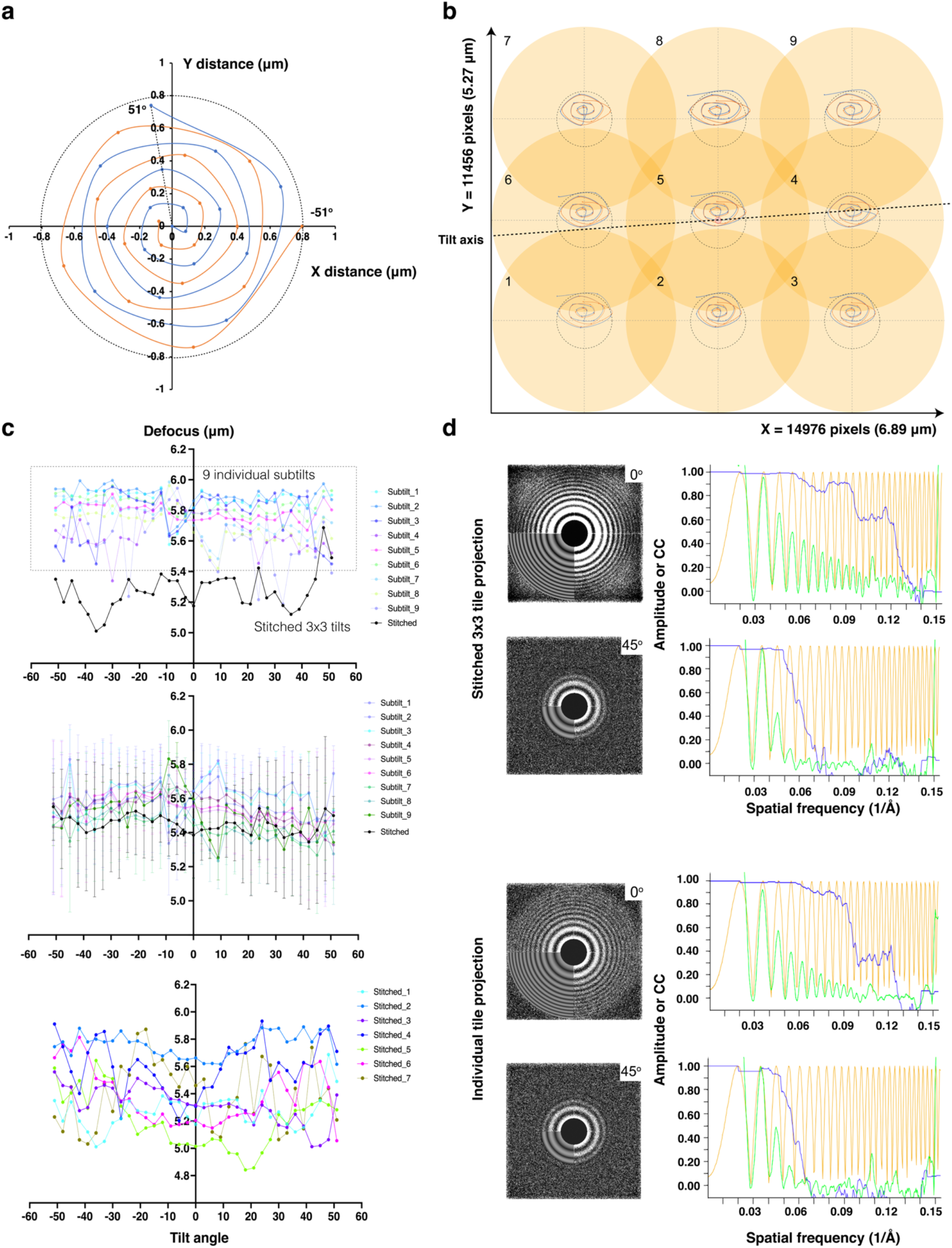
Defocus determination and CTF estimation of full stitched montage and individual tile tilt series. a. Benchmark spiral translation pattern applied to distribute electron dose. To introduce global translation offsets, the central tile of the montage follows spiral paths shifting outwards over the full course of the tilt series from -51° (orange points) to 51° (blue points) in a dose-symmetric, three-degree increment and group of 3 scheme. The translation maximum is 0.8 *µm* (black dotted circle, A_final_ = 1.5, Period = 3, Revolution = 15 as spiral shape parameters). **b**. Cross-correlation alignment of cosine-stretched image shifts at each tilt angle relative to 0° tilt in individual tile tilt series of the regular 3×3 montage pattern, using Tiltxcorr in IMOD. The image shift between each tilt follows a stretched spiral along the X axis when the optional scaling parameter of the translation is not applied. The image shifts of individual tile tilt series were overall within the maximum distance of 0.8 *µm* (black dotted circle). Numbers indicate the individual tile positions applied to both **(c)** and **(d). c**. Defocus values of nine individual tile sub-tilt series and the corresponding stitched 3×3 montage tilt series (pixel size of 4.603 Å), following the same global spiral shift in **(a, b)** were estimated using CTFPLOTTER in IMOD: one representative 3×3 stitched montage and nine individual tile tilt series (top), median variation at each tilt angle over multiple 3×3 montage and corresponding tile tilt series (n = 3, middle), and 7 stitched montage tilt series (bottom) are shown. A target defocus of -5 *µm* was applied by performing autofocusing prior to the tile collection at each tilt angle, along the tilt axis, 7.5 *µm* away from the center of the montage tile to account for the size of the montage (6.89 × 5.27 *µm*) and additional translation shift (0.8 *µm*). **d**. CTF estimation in CTFFIND4 by the 2D power spectra (left) from projections at 0°, 45°, and -45° from a representative 3×3 stitched montage tilt series and one of the individual tile tilt series (tile position 2 in **b**), and corresponding fitted 1D models (right) showing radially averaged amplitude spectrum (green), CTF fit (orange), and CTF fit score (blue).

**Supplementary Figure 10.**
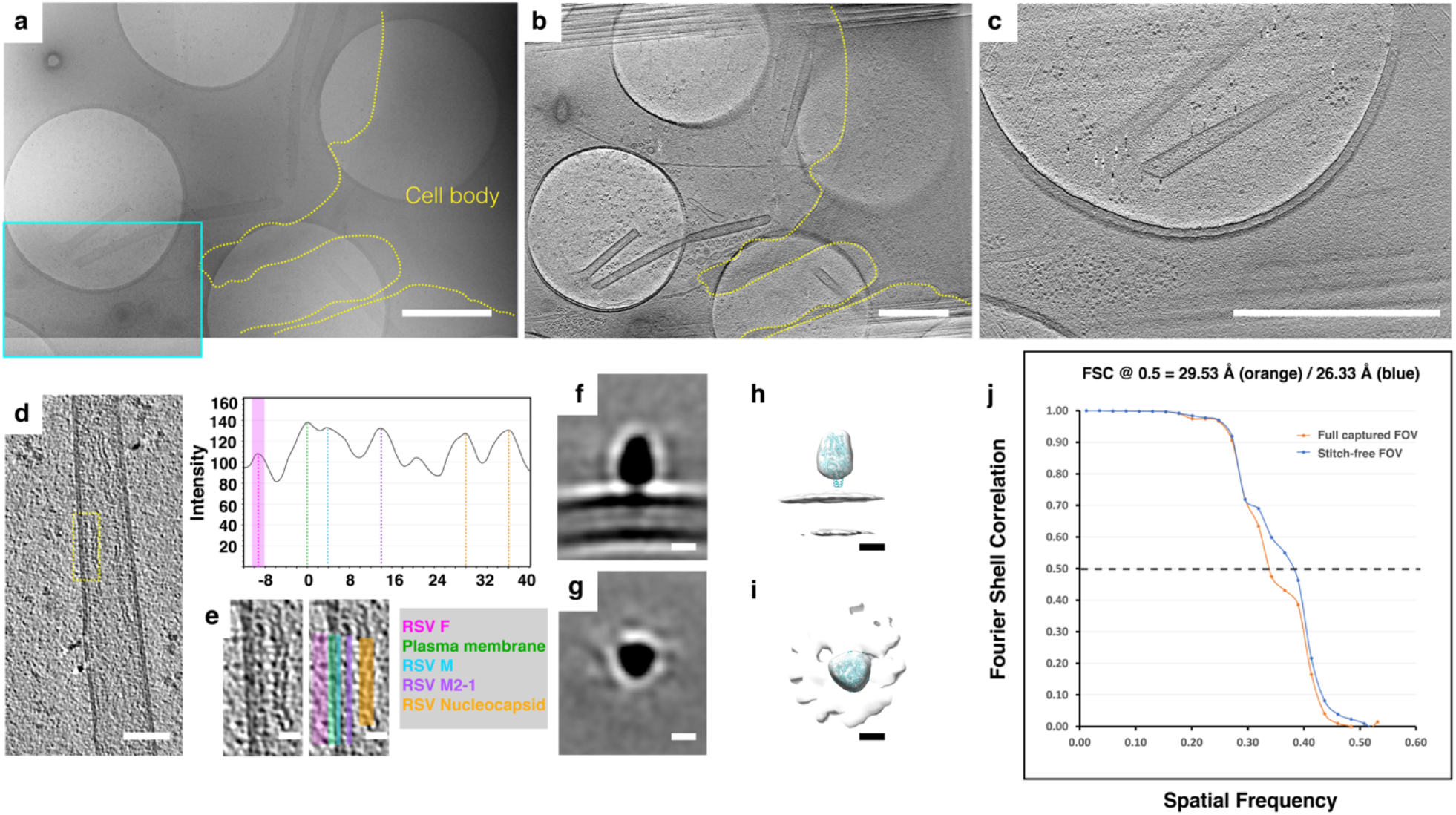
Sub-tomogram averaging of individual tile tomograms. **a**. 3×3 montage cryo-ET image tiles at 0° tilt of RSV-infected BEAS-2B cells. The cell boundary is delineated in yellow and one tile with viruses used for sub-tomogram averaging is highlighted in cyan. **b**. Central slice of the stitched 3×3 montage tomogram in **(a)**, cell boundary (yellow). **c**. Central ∼18 nm slice of the reconstructed tomogram (pixel size of 9.206 Å) of the tile highlighted in (**a**, cyan box). Scale bars, 1 *µm* in **a-c. d**. Reoriented Z-projection (9 nm thick) of the virion from the tomogram in **(c). e**. An enlarged view of virus (yellow dashed box in **d**) with virion components noted in different colors and a linear density profile across the virion region. Scale bars, 100 nm in **(d)**, 20 nm in **(e). f-i**. Sub-tomogram averages of RSV F from individual tile tomograms collected through the 3×3 montage cryo-ET workflow. The particles in overlapped stitched zones (15% in X and 10% in Y) were excluded. **f-g**. Slices from a sub-tomogram average of binned 2 individual tile tomograms (pixel size of 9.206 Å) of RSV F filtered to 27 Å in the side (**f**) and top (**g**) views. **h-i**. RSV pre-fusion trimer model (PDB: 4JHW, cyan) fit into an isosurface of the sub-tomogram average of F. Scale bars, 5 nm. **j**. FSC curves for sub-tomogram averages of F particles picked from full captured field of view (orange, C3 symmetry expansion, n = 23250) and field of view free from stitching overlaps (blue, C3 symmetry expansion, n = 20707) as stitch-free particles in sub-tilt tomograms.

**Supplementary Table 1.** Dose accumulation in e^-^/Å^2^/voxel at specified voxels in four sampling areas (radius of 10 × 10 × 1 voxel^2^ pivoting around the central sampling points 1 to 4, X and Y voxel positions specified, indicated in **Supplementary Fig. 4**) of six different collection schemes with the same 3×3 montage tile pattern using *TomoGrapher* simulation. The mean and standard deviation were calculated. The Tilt Strategy parameters is also specified in Supplementary Fig. 4 (3×3 tile pattern, pixel size of 4.603 Å, beam size of 3.1 *µm*, from -51° to 51° tilts with a 3° increment, 1 or 0.7 e^-^/Å^2^ per tile per tilt).

|  | Position 1<br>(x/y, 38/101) | Position 2<br>(x/y, 45/87) | Position 3<br>(x/y, 59/103) | Position 4<br>(x/y, 58/87) |
| --- | --- | --- | --- | --- |
| No translation (a) | 39.2 ± 5.2 | 85.1 ± 18.1 | 64.5 ± 7.7 | 105.1 ± 18.5 |
| No translation lower dose by 30% (b) | 27.5 ± 3.6 | 59.6 ± 12.5 | 45.2 ± 5.3 | 73.6 ± 12.9 |
| Translation (c) | 45.5 ± 3.2 | 77.1 ± 4.8 | 56.8 ± 7.4 | 79.2 ± 10.1 |
| Translation lower dose by 30% (d) | 31.8 ± 2.2 | 48.2 ± 3.3 | 39.8 ± 5.2 | 56.1 ± 7.1 |
| Translation with x-axial correction (e) | 45.2 ± 4.8 | 83.7 ± 10.8 | 66.7 ± 7.6 | 94.7 ± 10.8 |
| Translation with x-axial correction lower dose by 30% (f) | 31.7 ± 3.4 | 51.2 ± 7.6 | 46.7 ± 5.3 | 66.3 ± 7.5 |

